# De novo designed cyclic MC4R peptide agonist reduces food intake in mice

**DOI:** 10.64898/2026.05.19.721857

**Authors:** Victor Møller, Julie Maria Johansen, Randi Bonke Mikkelsen, Poanna Tran, Ashref Kayed, Nina Buch-Månson, Timothy P. Jenkins, Louise S. Dalbøge, Jens Christian Nielsen, Mads Mørup Nygaard

## Abstract

Deep learning-based structure prediction enables the design of peptide ligands without relying on naturally occurring scaffolds. However, most computationally generated peptides are not advanced beyond initial activity measurements, leaving the path to drug-like optimization and in vivo validation underexplored. Here we establish an end-to-end workflow for de novo peptide agonist discovery and maturation using the melanocortin-4 receptor (MC4R) as a model target. Using an AlphaFold2-based hallucination protocol implemented in ColabDesign, we generated more than 5,000 linear and head-to-tail cyclic candidate peptides directed towards the MC4R orthosteric pocket. Functional screening of a prioritized subset revealed measurable activity in 74% of linear peptides and 23% of cyclic peptides, from which we identified a cyclic agonist with an EC_50_ of 340 nM despite lacking the canonical melanocortin activation motif. We then performed systematic in vitro maturation by deep mutational scanning, half-life extender conjugation scanning, and a combinatorial optimization library, coupled with data-driven analysis to map sequence-activity relationships. These experiments identified an alternative activation motif centered on an APWR segment and yielded single-site variants with substantially improved potency. The most effective substitution, a proline at position 5, produced the E5P variant with an EC_50_ of 6.7 nM against the human melanocortin-4 receptor (hMC4R). Finally, central administration of E5P (10 nmol) reduced acute food intake in mice, providing in vivo proof of concept. Together, our results demonstrate a generalizable design-to-validation strategy for converting de novo peptide designs into optimized, pharmacologically active peptides, and expand the space of MC4R agonist chemotypes beyond endogenous melanocortins.

## Main

Peptide therapeutics occupy a unique position between small molecules and biologics, combining high target selectivity and efficacy with favourable safety profiles. However, their translation often depends on extensive optimization to improve stability, exposure and other drug-like properties^1–3^. Most peptide discovery campaigns begin with naturally occurring peptide ligands followed by iterative modification through substitution, cyclisation, or lipid conjugation^1–3^. Although effective, this strategy ties discovery to targets with suitable natural starting points and largely confines exploration to sequence and structural space near evolved scaffolds^3^.

Display-based technologies, including phage display, mRNA display and DNA-encoded libraries, have expanded access to non-natural peptide binders^4,5^. Despite their power, display-based discovery approaches impose challenges: enriched binders can be non-specific or selection-driven artefacts, and many require substantial downstream optimization before they show robust functional activity^5–10^. Recent advances in deep-learning-based structure prediction, particularly AlphaFold2, have opened an alternative route by enabling peptides to be designed directly against target structures^11–20^. Hallucination- and diffusion-based approaches have generated de novo binders and functional modulators^21–29^, but most studies remain focused on initial in vitro activity.

Structured experimental strategies can refine potency, stability, and pharmacokinetic properties. This can be obtained through iterative in vitro optimization, guided by systematic structure-activity relationship (SAR) analysis, which has proven efficient for advancing peptide therapeutics^30,31^. Although a few recent reports have extended validation of de novo designs into in vivo settings^26,29,32^, it remains under-explored whether de novo peptides can be systematically matured into pharmacologically useful ligands with in vivo efficacy.

This is especially relevant for G protein-coupled receptors (GPCRs), which account for ∼36% of approved drugs^33^, and where emerging work has begun to extend de novo design from target engagement to functional activity^34,35^. The melanocortin-4 receptor (MC4R), a class A GPCR central to energy homeostasis and body weight regulation^36^, is an established therapeutic target for obesity. However, endogenous melanocortin peptides activate multiple melanocortin receptors through a conserved HFRW motif^37,38^, complicating efforts to achieve subtype selectivity and motivating the search for alternative chemotypes. Design strategies that are not anchored to endogenous melanocortin scaffolds may therefore expand the accessible landscape of MC4R ligands and reveal distinct receptor interaction modes.

Here we combine AlphaFold2-based hallucination, using ColabDesign, with structured experimental maturation to discover de novo peptide agonists of human MC4R (hMC4R). We designed linear and head-to-tail cyclic peptides targeted to the orthosteric pocket land refined this scaffold through deep mutational scanning, half-life extender conjugation scanning and combinatorial optimization, supported by data-driven analysis of sequence-activity relationships. We further characterized receptor selectivity across the melanocortin receptor family and evaluated in vivo activity by measuring food intake following central administration in mice. Together, we establish a design-to-validation framework for converting de novo peptide designs into optimized, in vivo-active ligands.

### De novo peptides with noncanonical motif

To test whether AF2-guided hallucination could generate functional ligands for MC4R, we used ColabDesign and AFCycDesign to design de novo peptides targeted to the hMC4R orthosteric pocket^15,25^ (Fig. 1a). We selected residues lining the pocket as design hotspots to bias designs toward receptor engagement and potential functional activity. In total, approximately 5,000 peptide sequences were generated in silico, spanning multiple lengths and comprising both linear and head-to-tail cyclic peptides to systematically explore conformational space and enhance structural diversity. We prioritized predicted receptor-peptide complexes using two AF2 confidence metrics, the predicted local distance difference test (pLDDT) and the interface predicted template modelling score (ipTM), by integrating them into a composite score. Such heuristic composite metrics derived from AF2 are widely used to rank and triage de novo designs^13,15,22^. Sequences were ranked according to this composite confidence score, and top-scoring candidates were selected for synthesis to determine whether they modulated hMC4R signaling in cells (Fig. 1b,c).

**Fig. 1:**
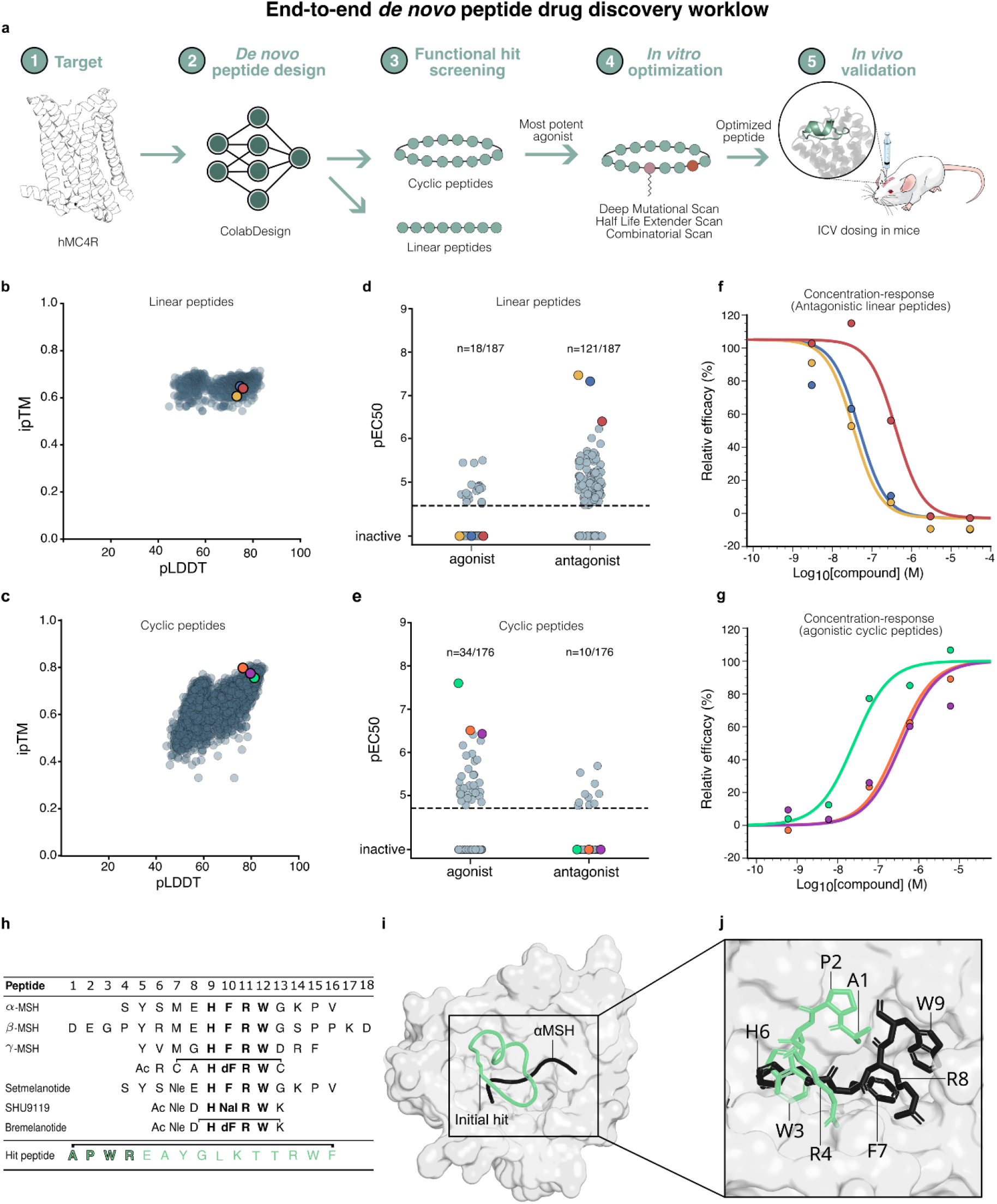
Workflow and distributions of linear and cyclic de novo peptide designs to hMC4R. **a**) De novo design workflow of linear and cyclic peptides targeting hMC4R, followed by in vitro maturation of top cyclic hit, finalized with in vivo validation of optimized peptide by measuring food intake in intracerebroventricular-dosed mice. In vitro maturation included various sequence library designs; deep mutational scan (DMS), half-life extender (HLE) scan and a follow-up library combining the beneficial substitutions and HLE-conjugations in previous designs. Peptides were synthesized with solid-phase peptide synthesis (SPPS) and in vitro assays determined features including receptor potency. **b)** In silico structural confidence metrics (pLDDT and ipTM of 1200 linear de novo peptides designed to bind to hMC4R. **c)** In silico structural confidence metrics (pLDDT and ipTM) of 3803 cyclic de novo peptides designed to bind to hMC4R. **d)** Top 189 linear designs were selected and 187 synthesized and evaluated as crude peptides, revealing agonist and antagonist pEC_50_ distribution with a hit rate of 74% in the linear de novo hit library. **e)** Top 184 cyclic designs were selected and 176 synthesized. Agonist and antagonist pEC_50_ distribution of crude peptides demonstrate a hit rate of 23% in the cyclic de novo hit library. **f)** Experimental concentration-response curves at hMC4R in antagonist mode of top 3 linear hits as crude. **g)** Experimental concentration-response curves at hMC4R in agonist mode of top 3 cyclic hits as crude. **h)** Sequences of known MC4R peptide ligands. Nal denotes (2R)-2-amino-3-(naphthalen-2-yl)-propanoic acid, a non-canonical tryptophan-like aromatic residue. **i)** Structural superposition of the hit peptide (green) and α-MSH (black) bound to MC4R (grey). The predicted hit-hMC4R complex generated with ColabDesign was aligned to the experimental mouse MC4R (mMC4R) structure (PDB 7F53) based on receptor backbone atoms. **j)** The hit peptide and α-MSH shown as cartoon structures zoomed in on the receptor pocket. Binding motifs: position 1-4 (APWR) of the best de novo hit peptide and position 6-9 (HFRW) of α-MSH in MC4R pocket shown as sticks.

We synthesized prioritized designs as crudes by solid-phase peptide synthesis (SPPS) and screened them for hMC4R agonist and antagonist activity in cell-based cAMP assays. For cyclic peptides, we restricted designs to 13 and 16 residues and filtered sequences to ensure compatibility with head-to-tail cyclisation, yielding 184 cyclic peptides of which 176 were successfully synthesized. In parallel, we selected 189 linear peptides using the same confidence criteria. In the functional screening, 74% of linear peptides and 23% of cyclic peptides showed measurable agonist or antagonist activity. Linear designs more frequently acted as antagonists, whereas cyclic peptides were enriched for agonist activity (Fig. 1d,e). This was also observed in individual concentration-response curves for all peptides including the three most potent linear peptide hits showing antagonist behaviour (Fig. 1f), and the three most potent cyclic hits showing agonist behaviour (Fig. 1g). We selected the most potent cyclic agonist for follow-up. Upon purification, it retained full efficacy with an EC_50_ of 340 nM (Table S1), and head-to-tail cyclic peptides provide a tractable starting point for drug-like optimization. The lack of free termini can increase resistance to exoproteases and may improve proteolytic stability and in vivo exposure^2,39^; with further optimization, such scaffolds may also support improved permeability and, in turn, the possibility of oral delivery^40^. Together, this provides a de novo starting point for further optimization of the hMC4R agonist.

Because the best agonist emerged from designs unconstrained by endogenous melanocortin sequences, we examined how the active de novo peptides related to the established melanocortin pharmacophore. AF2 modelling suggested that all the active peptides bound within the hMC4R orthosteric pocket but adopted diverse backbone conformations. Linear peptides typically inserted one terminus deeply into the pocket, whereas cyclic peptides tended to occupy a larger fraction of the pocket volume (Fig. S1). Notably, none of the de novo peptides contained the conserved HFRW activation motif characteristic of endogenous melanocortins^37,38^, as illustrated by known ligand alignments (Fig. 1h) and the mouse MC4R:α-MSH complex (Fig. 1i,j). Instead, motif analysis of the cyclic hit library revealed a 16-fold enrichment of a reversed WR sub-motif (Fig. S2), suggesting a distinct interaction pattern within the orthosteric pocket.

To connect sequence-level enrichment to putative binding modes, we clustered the functionally active cyclic peptides by contact fingerprints to the hMC4R and observed multiple distinct binding clusters within the same orthosteric pocket (Fig. S3). Structural comparison of the hit peptide to α-MSH further suggested a different backbone orientation in the pocket (Fig. 1i) and a shift in how the APWR segment was positioned relative to the HFRW motif of α-MSH (Fig. 1j). Together, these observations suggest that de novo design can yield hMC4R agonists compatible with an alternative activation motif, motivating systematic in vitro maturation to optimize pharmacological properties through targeted sequence and chemical modifications.

### In vitro maturation defines agonism

To convert the cyclic de novo hMC4R agonist (head-to-tail cyclized APWREAYGLKTTNRWF) into a more potent peptide with drug-like properties, we performed a multi-stage in vitro optimization campaign. We combined deep mutational scanning (DMS) to map sequence-activity relationships, half-life extender (HLE) conjugation scanning to probe positional tolerance to lipidation, and a combinatorial library to test whether beneficial substitutions could compensate for potency losses associated with HLEs. All variants were synthesized and screened for hMC4R agonist and antagonist activity. We initially normalized potencies to the hit peptide in the DMS and HLE scan libraries, and to the W15K_ε_C18DA variant in the combinatorial library (Fig. 2a). Then we corrected for batch effects between libraries not affecting the rank ordering between crude and purified peptides (Fig. S4, Fig. S5). We first used the DMS to identify which positions modulated potency of the initial cyclic hit peptide.

**Fig. 2:**
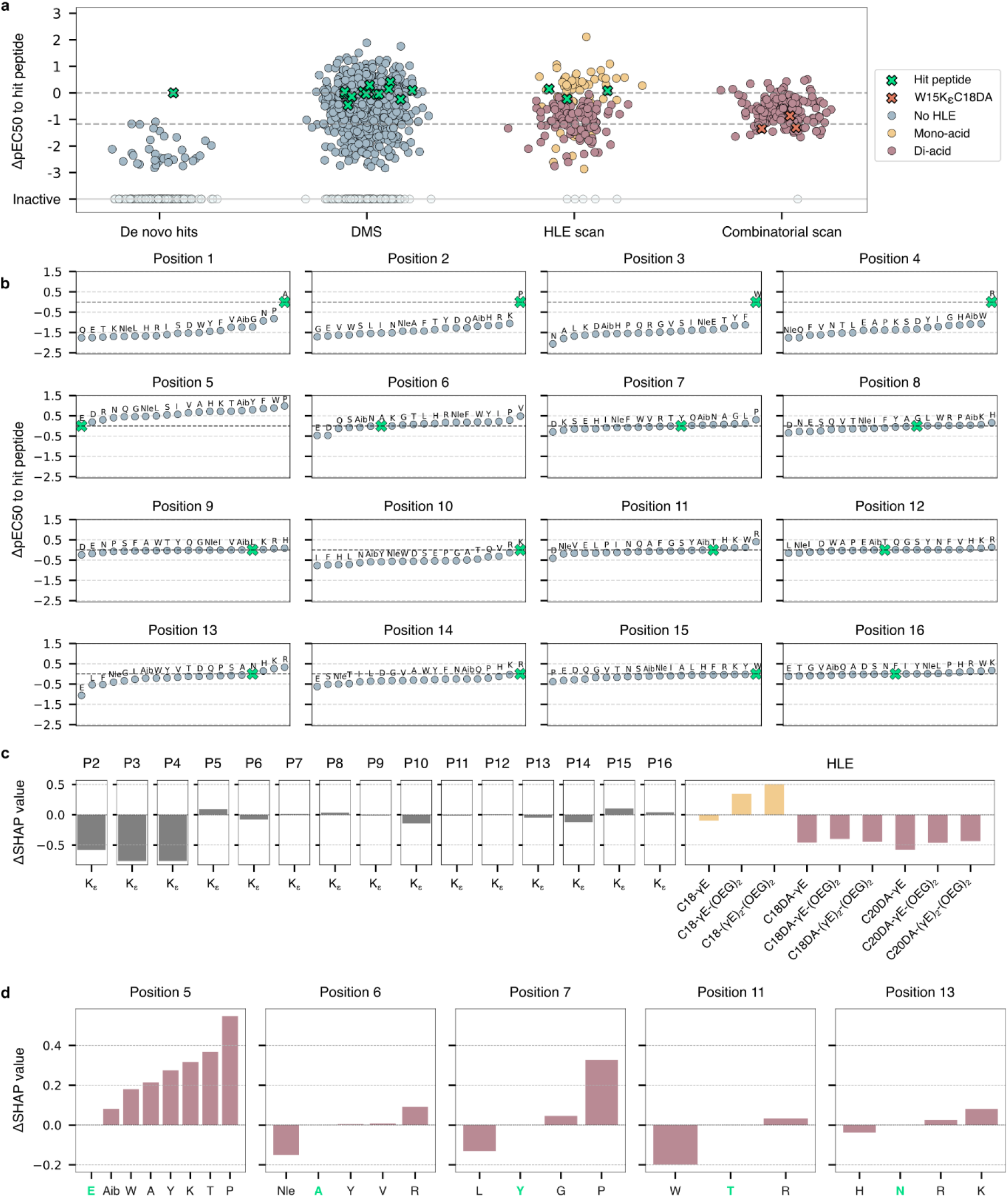
In vitro measurements and ML model predictions of de novo hits, DMS, HLE scan and combinational scan. **a**) Relative agonist pEC_50_ change of the peptide variants in each library relative to the best hit peptide. Upper grey dashed line at y=0 indicates hit peptide potency, lower grey dashed line indicates average W15KεC18DA variant potency. Green crosses are replicates of best hit peptides, orange crosses are the W15KεC18DA variant, light blue datapoints indicate no conjugation of an HLE to the peptide variants, yellow datapoints indicate attachment of a mono-acid, while purple datapoints indicate attachment of a di-acid. Grey transparent datapoints in the bottom highlight inactive peptides. **b**) Random forest regression model predictions of potency according to position and residue trained on DMS data. Highlighted in green are the amino acids corresponding to the hit peptide sequence. pEC50 values are relative to compound the hit peptide. **c**) SHAP values from random forest regression trained on the HLE scan data. Positive values correspond to a positive effect on receptor potency while a negative SHAP value indicate a negative contribution to receptor potency in relation to the hit peptide. The hit peptide was conjugated with 9 different mono- or di-acids with varying linker composition, L-γ-glutamyl (γE) and 3,8-dioxa-aminooctanoic acid (OEG) via the sidechain of a lysine substitution (Kε). Mono-acids are coloured in yellow and di-acids in pink. **d**) SHAP values from random forest regression trained on the combinatorial library design. The sequence of the hit peptide conjugated with a C18DA-γE-(OEG)_2_ di-acid at position 15 (W15K_ε_C18DA) is highlighted in green on the x-axis and correspond to SHAP value of 0.

Across the DMS library (835 variants comprising single-, double-, and triple-substitution mutants), most substitutions preserved agonist behaviour, with relatively few variants converting to antagonists (Fig. 2a, Fig. S6) and most substitutions did not cause an increase in fibril formation or turbidity (Fig. S7). Despite this overall functional robustness, potency effects spanned more than five orders of magnitude, with the most favourable substitutions improving potency by up to ∼100-fold relative to the hit peptide (Fig. 2a). Random-forest regression trained on the DMS dataset identified positions 5, 6, 7, 11, and 13 as major contributors to potency modulation (Fig. 2b). In contrast, substitutions at positions 1-4 consistently drastically reduced activity (Fig. 2b). Structural modelling supported this functional constraint, showing that residues 1-4 engage the deepest region of the MC4R binding pocket (Fig. 1i,j), defining an APWR motif critical for receptor activation. Having established the sequence constraints governing receptor activation, we next examined which positions accommodated HLE conjugation to support pharmacokinetic optimization.

Conjugations with HLEs are commonly used to prolong half-life via reversible serum albumin binding thereby decreasing renal elimination and susceptibility to enzymatic degradation^41^. To explore pharmacokinetic optimization without affecting potency, we performed a systematic HLE conjugation scan comprising 173 variants in which HLEs were attached via lysine side chains at positions 2-16. We evaluated various octadecanoic (C18), octadecanedioic (C18DA), eicosanoic (C20) and eicosanedioic (C20DA) acids with various linker moieties. For simplicity, C18- and C20-based HLEs are referred to as mono-acids, whereas C18DA- and C20DA-based HLEs are referred to as di-acids.

Most HLE-conjugated variants retained agonist activity and did not convert to antagonists (Fig. 2a, Fig. S8) while fibril formation and increased turbidity were observed for some di-acid extenders (Fig. S9). Mono-acid conjugation occasionally improved potency, whereas di-acid conjugation generally reduced activity (Fig. 2a). SHAP analysis of a random-forest model trained on the HLE dataset revealed strong positional effects, with conjugation at positions 2-4 being highly detrimental, and positions 5 and 15 showing greater tolerance (Fig. 2c). Across linker composition, mono-acid conjugates exhibited positive SHAP values, whereas all di-acid conjugates contributed negatively to potency (Fig. 2c). These results suggested that achieving potent di-acid conjugates would likely require compensatory sequence changes.

Finally, to assess whether favourable substitutions could compensate for the potency penalty associated with di-acid conjugation, we designed a combinatorial optimization library centred on a position-15 di-acid conjugate. Here we systematically combined the di-acid with beneficial substitutions identified in the DMS at positions 5, 6, 7, 11, and 13 (181 variants). The peptides retained agonist activity without detectable antagonism (Fig. S10). Fibril formation increased at pH 4, whereas turbidity increased for some peptides at pH 7 but not at pH 4 (Fig. S11). This library revealed strong context dependence: proline or tyrosine substitutions at position 5, and proline at position 7, substantially improved potency in the presence of di-acid conjugation (Fig. 2d). In contrast, simultaneous proline substitutions at both positions did not yield additive gains, consistent with non-linear epistatic interactions between backbone rigidification sites^42–45^ (Fig. S12). Collectively, these data defined a set of substitutions and conjugation sites for confirmation in purified compounds and evaluation of receptor-family selectivity.

### Altered subtype selectivity of compounds

Although crude peptides enabled rapid, cost-efficient screening and optimization^46^, the resulting receptor potency estimates were derived from non-purified samples and single-replicate 5-point concentration series which are best interpreted as relative rankings rather than definitive EC_50_ values. We therefore synthesized selected peptides at high purity and performed 11-point concentration series for detailed pharmacological characterization, including melanocortin receptor-family profiling (Fig. 3, Table S1). We focused our analysis on the endogenous MC4R ligand α-MSH, the initial de novo hit, the E5P variant and the E5KεC18:Y7P variant (Fig. 3a).

**Fig. 3:**
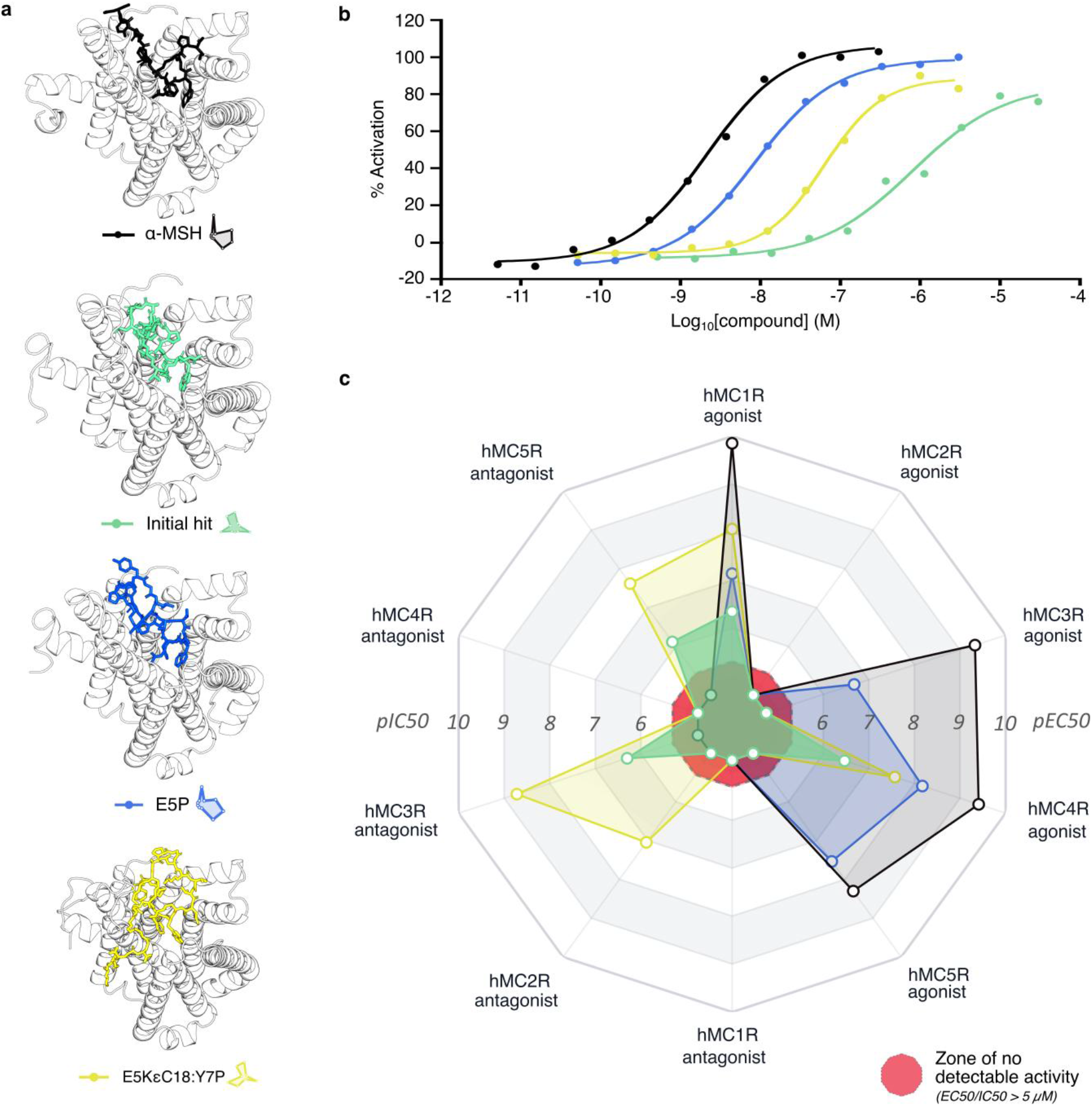
Selectivity profiling across the melanocortin receptor family. **a)** Predicted structures of α-MSH, the initial hit, variant E5P and variant E5KεC18:Y7P in context of MC4R. Structures are predicted with Boltz-2^47^. Low-confidence disordered region of the MC4R N-terminal is not shown. **b)** Experimental concentration-response curves at hMC4R in agonist mode of selected purified peptides. The shown curves are representative data from one of the independent experiments contributing to the reported average potency values. **c)** Radar plot summarizing agonist and antagonist potencies of α-MSH and purified de novo hMC4R agonist variants across hMC1R-hMC5R (colours as indicated). Each spike corresponds to a receptor measured in agonist mode (pEC_50_) or antagonist mode (pIC_50_), as labelled; higher values indicate higher potency. The central red region denotes no detectable activity at the highest concentration tested. Measurements outside the assay window are reported as “>“ in Table 2 (α-MSH: EC_50_/IC_50_ > 5,000 nM; de novo peptides: EC_50_/IC_50_ > 50,000 nM) and are plotted at the same minimum radius to indicate inactivity despite different assay ceilings.

The purified hit peptide exhibited an hMC4R agonist potency of 340 nM (Fig. 3b, Table S1). Single-site substitutions identified in the optimization campaign translated robustly upon purification. In particular, introducing a proline at position 5 (E5P) improved potency by ∼51-fold to an EC_50_ of 6.7 nM (Table S1, Fig. 3b), representing the strongest single substitution across all libraries. A proline substitution at position 7 (Y7P) also improved potency, albeit slightly more modestly (EC_50_ = 39 nM; Table S1).

We next characterized selected conjugated variants in follow-up in vitro assays. Mono-acid conjugation enhanced potency in selected sequence contexts, whereas di-acid conjugation generally reduced agonist activity; however, several di-acid conjugates remained active in the nanomolar range when combined with favourable substitutions (Table S1). Given the utility of di-acids for half-life prolongation^41^, we explored whether introducing the di-acid via a 4-aminoproline (Pγ; Fig. S13) rather than a lysine side chain (Kε) could recover potency in di-acid-modified variants. We did not observe substantial differences between compounds carrying combined modifications and those with separate substitutions and HLE conjugations. Linearized versions of the hit peptide and optimized variants were inactive (Table S1), indicating that head-to-tail cyclization was required for hMC4R agonism in this chemotype. With purified peptides in hand, we next quantified off-target activity across the melanocortin receptor family.

To benchmark selectivity, we profiled α-MSH and purified de novo variants across hMC1R, hMC3R, hMC4R and hMC5R in agonist and antagonist assay formats (Fig. 3c, Table S1). As expected, α-MSH showed broad agonism, with sub-nanomolar potency at hMC1R, hMC3R, and hMC4R (EC_50_ = 0.14-0.47 nM) and low-nanomolar potency at hMC5R (EC_50_ = 5 nM), while no activity was detected at hMC2R within the tested concentration range, in agreement with previously reported receptor selectivity and potency profiles^36^.

Across this panel, none of the de novo variants were exclusively hMC4R-selective. Instead, each compound exhibited a receptor- and modification-dependent pattern of agonism and/or antagonism that differed from α-MSH. The original hit peptide activated hMC4R with EC_50_ = 340 nM and showed partial agonism at hMC1R (EC_50_ = 450 nM; 61% efficacy), with no detectable agonism at hMC2R, hMC3R, or hMC5R. In this panel, the hit peptide instead antagonized hMC3R and hMC5R (IC_50_ = 500 nM and 760 nM, respectively).

The E5P variant, with hMC4R agonist potency of 6.7 nM, displayed measurable agonism at hMC1R (73 nM) and hMC5R (29 nM) and weak partial agonism at hMC3R (210 nM; 38% efficacy), while no agonism was detected at hMC2R (>50 µM). These values correspond to ∼11-fold (hMC1R), ∼4-fold (hMC5R), and ∼31-fold (hMC3R) weaker agonist potency relative to hMC4R, with the hMC3R response additionally limited in efficacy. By contrast, addition of HLE conjugation often shifted cross-reactivity toward antagonist activity at non-hMC4R receptors (Fig. 3c; E5KεC18:Y7P, yellow trace), a trend that was observed across multiple HLE-conjugated compounds (Table S1). W15KεC18 and Y7P:W15KεC18 maintained hMC4R agonism (EC_50_ = 73 nM and 55 nM), showed partial agonism at hMC1R, but antagonized hMC2R, hMC3R, and hMC5R (IC_50_ = 41-250 nM). The combined HLE-conjugated variant E5KεC18:Y7P showed hMC4R agonism (EC_50_ = 27 nM) together with potent hMC1R agonism (EC_50_ = 8.6 nM), while acting as a strong antagonist at hMC3R (IC_50_ = 1.9 nM), hMC5R (IC_50_ = 24 nM), and hMC2R (IC_50_ = 90 nM). Together, these data established E5P as a potent agonist with its strongest agonist potency at hMC4R within the receptor-family panel and motivated an in vivo test of central MC4R engagement.

### Central dosing reduces food intake

Given the encouraging in vitro potency and minimal sequence modification, we selected the E5P variant for in vivo evaluation. Lean male NMRI/RjHan mice were fasted 6 hours prior to dosing and received a single intracerebroventricular (ICV) administration of E5P (10 nmol) or vehicle (n=8/group) immediately prior to the onset of the dark phase. Animals were randomized based on body weight and 24-hour food intake prior to dosing (Table S2). Food and water intake were continuously monitored for 24 hours using a comprehensive metabolic phenotyping system (PhenoMaster) (Fig. S14).

As expected, the most noticeable differences in food and water intake were observed during the dark phase of the light-dark cycle, corresponding to the first 12 hours post-dosing, when food intake is typically highest in mice. Relative to vehicle-treated controls, E5P produced a rapid and pronounced reduction in cumulative food intake, with an approximate 34% decrease within the first hour and a 23% reduction at 4 hours post-dose (Fig. 4a, Table S3). Although the anorectic effect diminished over time, cumulative food intake remained significantly reduced at 8 hours post-administration (Fig. 4b), representing the largest sustained difference between groups. No significant differences in food intake were detected beyond this time point.

**Fig. 4:**
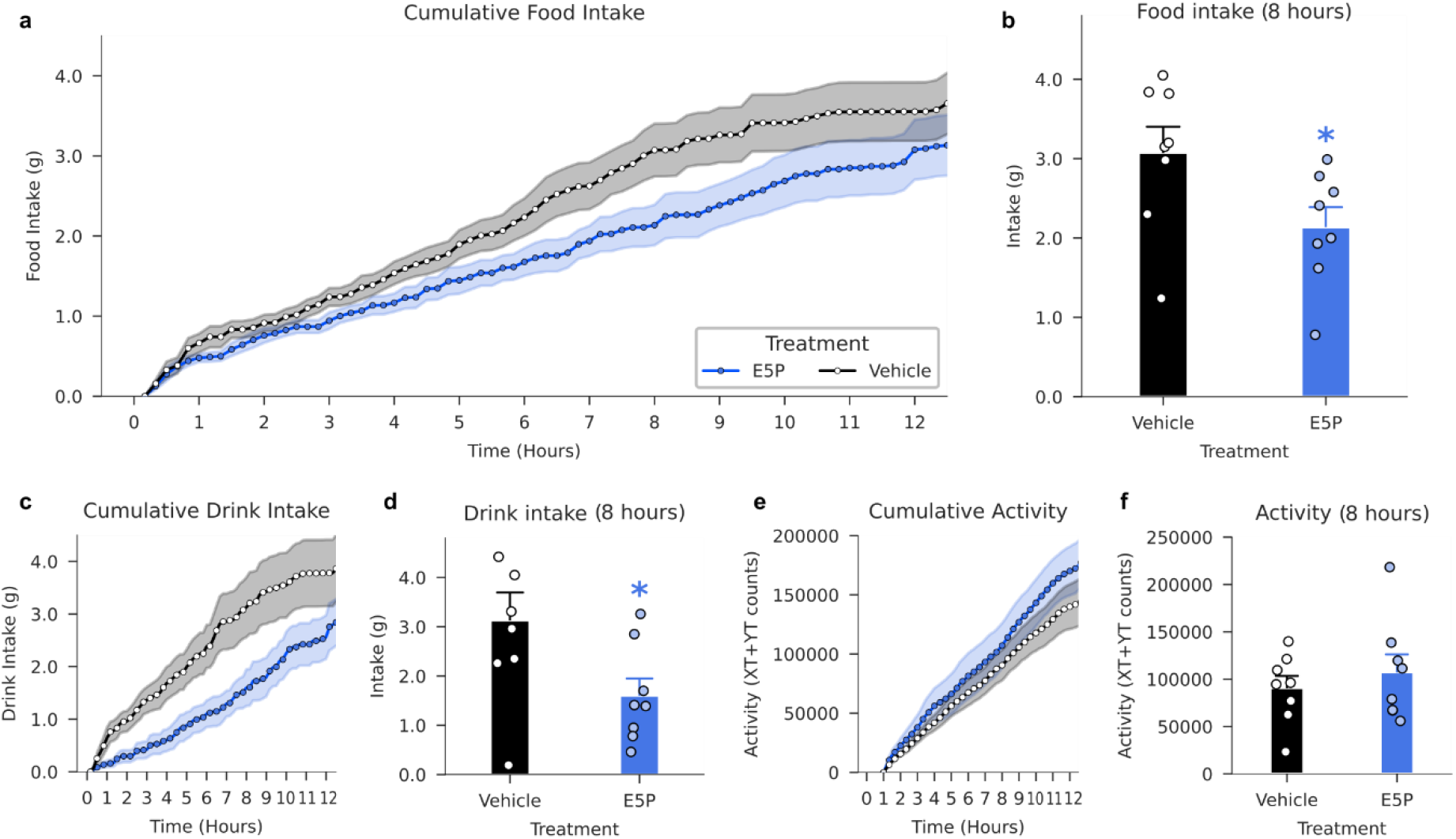
Intake and activity of ICV-dosed animals between vehicle-treated (white) and E5P-treated (blue) mice. **a**) Cumulative food intake over the first 12 hours post-dosing, corresponding to the dark phase, when mice exhibit peak feeding activity. **b**) Cumulative food intake from 0-8 hours. **c**) Cumulative water intake over the first 12 hours. **d**) Cumulative water intake from 0-8 hours. **e**) Cumulative activity data over the first 12 hours measured in XT+YT counts corresponding to the sum of infrared beam breaks in the X- and Y-axes recorded by the PhenoMaster system. **f**) Cumulative activity data from 0-8 hours. Values are expressed as mean of n=8 ± SEM. Two-tailed unpaired t-test performed with * meaning p-value < 0.05 compared to vehicle group.

Water intake followed a similar temporal pattern, with a marked reduction during the early post-dose period and gradual recovery thereafter (Fig. 4c,d). Furthermore, E5P treatment did not influence locomotor activity (Fig. 4e,f), indicating that the observed reductions in food and water intake were not secondary to decreased activity. Together, these data demonstrate proof-of-concept of a de novo designed and experimentally optimized MC4R agonist peptide with relevant in vivo activity.

## Discussion

This study provides direct evidence that generative protein design can produce entirely new receptor-modulating chemotypes that function in vivo. By linking structure-guided peptide generation with systematic experimental maturation and physiological validation, we show that computational design can extend beyond target engagement toward pharmacological activity in living systems. Using MC4R agonism as a model system, we establish a design-to-validation framework that connects de novo peptide generation to functional refinement and in vivo proof-of-concept.

Recent deep learning-enabled design strategies have demonstrated that de novo designed proteins and peptides can be engineered to engage biological targets with high structural precision. However, most efforts have focused on generating binders or functional activity in vitro, leaving unresolved whether such computationally created molecules can be matured into pharmacologically active ligands capable of modulating physiological systems. Our results bridge this gap by demonstrating that generative peptide design can be coupled to systematic functional optimization and translational validation within a single workflow. A notable finding is that the optimized agonist lacks the canonical melanocortin HFRW pharmacophore, instead relying on an alternative APWR-centered interaction motif. This observation indicates that generative design can access functional receptor interaction modes distinct from endogenous ligands. Expanding the accessible chemotype landscape beyond evolutionarily conserved pharmacophores may enable the discovery of receptor modulators with new selectivity profiles, signalling behaviours, and functional properties that are difficult to achieve through traditional ligand discovery approaches^48^.

Systematic experimental maturation proved critical for converting initial hit scaffold into potent receptor agonists with more than 50-fold improvement (E5P; EC_50_ 6.7 nM at hMC4R). Sequence-activity mapping defined the positions that constrained receptor activation and revealed non-additive effects among mutations, while half-life extender conjugation exposed strong position- and chemistry-dependent trade-offs in apparent potency. This approach demonstrates how de novo designed peptide scaffolds can be progressively refined through targeted functional feedback, providing a generalizable strategy for optimizing de novo biomolecules.

Although demonstrated here for MC4R, the workflow is receptor-agnostic and should be applicable to diverse membrane proteins and soluble targets where structural information is available. Generative design may be particularly valuable for receptors lacking well-defined endogenous ligands or for systems in which conserved pharmacophores constrain subtype selectivity. By enabling the creation of entirely synthetic ligands, this framework expands the accessible design space for therapeutic modulation of biologically important targets.

Within the melanocortin receptor family, endogenous ligands exhibit broad receptor activity due to highly conserved activation motifs, presenting intrinsic challenges for subtype-selective pharmacology. The alternative chemotype identified here alters subtype interaction profiles relative to the endogenous peptide ligands, illustrating how de novo design may provide complementary routes toward engineering receptor selectivity. Future integration of structure-guided modelling, subtype-specific functional assays, and targeted sequence diversification may further refine selective receptor activation using generative approaches. Several translational aspects remain to be explored.

While central administration established physiological proof-of-concept, systemic delivery strategies, pharmacokinetic optimization, and long-term efficacy studies will be necessary to evaluate therapeutic potential. Additionally, lipid conjugation influenced potency under albumin-depleted assay conditions^49^, and further work will be required to understand how protein binding and in vivo distribution affect pharmacodynamic properties. These challenges represent natural extensions of the framework rather than limitations of the design paradigm itself.

More broadly, this work illustrates how deep learning-based generative models are beginning to transition from tools for structural prediction to platforms for engineering functional biomolecules. As computational design becomes increasingly integrated with automated synthesis and high-throughput functional screening, the boundary between digital molecular design and experimental pharmacology may continue to narrow. The ability to create biologically active molecules from de novo sequences suggests a future in which peptides can be designed on demand for specific therapeutic objectives. Together, these findings establish a practical pathway from generative peptide design to in vivo functional validation and demonstrate that de novo peptide design can expand the landscape of receptor pharmacology beyond the constraints of natural occurring peptide ligands.

## Methods

### In Silico De Novo Peptide Design

Prior to the de novo designs, a crystal structure of hMC4R (PDB: 6W25) was selected as structural target template. This structure was selected because it offers a robust, internally consistent structure with well-defined local geometry at the binding site useful for de novo peptide generation and scoring. In addition, GPCRdb annotates 6W25 as an intermediate conformational state, which we considered a practical starting point to reduce bias toward a single end-state conformation and thereby increase the likelihood of identifying successful de novo binders^1,2^. To minimize the generation time, the target was cropped so the intracellular part of the receptor was disregarded. This resulted in a 157 amino acids long target structure.

AlphaFold2 binder hallucination was performed using ColabDesign’s setup with following settings listed below^15^. Furthermore, we guided the binder design towards the receptor pocket by selecting 3 hotspot amino acids located around the intended binding site. A subset of three residues was selected as hotspot from the following positions [97, 129, 197, 258, 261, 284, 288].

- model = mk_afdesign_model(protocol=“binder”)
- optimizer = pssm_semigreedy
- GD_method = sgd
- use_multimer = False
- target_flexible = False
- learning rate = 0.1
- dropout = True
- num_models = 5
- num_recycles = 0
- norm_seq_grad = True

Both linear and cyclic peptides were designed. For the cyclic peptide designs, we utilized the AfCycDesign implementation within the ColabDesign v1.1.0 framework, resulting in a head-to-tail cyclization of the designed peptides^25^. A total of around 1200 binders were designed for linear peptide of lengths 8, 10, 11, and 12 amino acids, while a total of around 3800 binders were designed as cyclic peptides with lengths 13 and 16 amino acids. We converted methionines into norleucine of the linear designs and avoided cysteines and methionines for the cyclic designs by running with the rm_aa = “C,M” setting due to potential challenges of side-chain reactivity during peptide synthesis^53^. Additionally, these cyclic designs were filtered based on sequences containing at least one alanine as the head-to-tail *N*-acyl-*N′*-methylacylurea (MeNbz) linker chemistry required an alanine for cyclization. The sequences were rotationally aligned to obtain an alanine at position 1.

We ranked the designs based on the structural confidence metrics by taking the product of the individual pLDDT and ipTM scores. Top-scoring 92 cyclic peptides of length 13 and 16 based on the abovementioned combined confidence score were selected for synthesis, resulting in a library of 184 cyclic peptides. In addition to the de novo designs, three replicate negative control peptides of lengths 13 and 16 were synthesized, comprising sequence-rearranged variants of the de novo designs. This yielded a cyclic peptide library comprising 190 designs, of which 182 were successfully synthesized; 176 of these corresponded to de novo designed cyclic peptides. In parallel, the top 189 linear peptide designs were selected using the same scoring criteria, of which 187 were successfully synthesized.

#### Boltz-2 predictions of structures

Complex structures used in Fig. 3 were predicted using Boltz-2^45^ with default inference settings. Multiple sequence alignments (MSAs) were supplied for hMC4R and for the endogenous ligand α-MSH. In contrast, de novo designed peptides were predicted without MSAs. For each complex, the top-ranked model according to Boltz internal confidence metrics was selected for visualization.

### Peptide library synthesis

Peptide library synthesis was performed using Fmoc-based solid-phase peptide synthesis (SPPS) on automated parallel peptide synthesizer, following the procedure described by Nielsen et al. (2024)^31^. Briefly, peptides were assembled using iterative cycles of Fmoc deprotection and amino acid coupling under heated conditions. HLE-conjugated peptides were generated via site-specific incorporation of protected lysine residues and subsequent on-resin lipidation as previously described.

Head-to-tail cyclization of selected peptides was achieved using *N*-acyl-*N′*-methylacylurea (MeNbz) linker chemistry. Linear peptides were synthesized on resin with incorporation of the 3,4-methyl-diaminobenzoic acid (MeDbz) linker at the C-terminus, followed by conversion to MeNbz using 5 equiv 4-nitrophenyl chloroformate and 25 equiv diisopropylethylamine, enabling intramolecular amide bond formation upon cyclization in phosphate buffer at pH 7. After cyclization, desulfurization was carried out to generate the desired alanine residue at the cyclization junction using 17 equiv sodium tetraethylborate and 5 equiv tris(2-carboxyethyl)phosphine hydrochloride, as described in Sun et al. (2022)^8^.

Following synthesis, peptides were cleaved from the resin, cyclized and desulfurized, and obtained as crude libraries. Crude peptide libraries were obtained without further purification and characterized by LC-MS. Selected peptides were purified by preparative reverse-phase HPLC as described by Nielsen et al. (2024)^31^.

### In Vitro Functional Assay for Melanocortin Receptor Potency

#### Cell lines

CHO-K1 cells were transiently transfected with hMC1R, hMC2R, hMC3R, hMC4R or hMC5R in pcDNA3.1(+)-N-DYK and frozen as assay-ready cells after 24 h. For MC2R, cells were co-transfected with hMRAP1α. All plasmids were obtained from GenScript Biotech (Rijswijk, Netherlands).

#### Assay

Frozen assay-ready cells overexpressing hMC1R, hMC2R/hMRAP1α, hMC3R, hMC4R or hMC5R were thawed, resuspended in DPBS with 0.05% casein, 0.5 mM IBMX and 1 mM CaCl_2_ and added as 4000 cells/well in a white 384-well microplate (Corning, cat. no. 4513). In agonist mode, the cells were then immediately stimulated for 30 min at room temperature with graded concentrations of test compounds. In antagonist mode, cells were stimulated with a fixed concentration (∼EC_90_) of α-MSH (Bachem, cat. no. 4008476) for hMCR1, hMCR3, hMCR4 and hMCR5 or ACTH(1-39) (Bachem, cat. no. 4030325) for hMC2R in the presence of graded concentrations of test compounds. Intracellular cAMP accumulation was measured using an HTRF Gs dynamic kit (Revvity, cat. no. 62AM4PEC), where the assay reagents were added as per the manufacturer’s instructions and time-resolved fluorescence energy transfer recorded after one hour on a CLARIOstar (BMG Labtech) plate reader.

Crude peptides were screened using 10-fold serial dilutions across 5 concentrations, n=1 time. Fresh pipette tips were used for the first two serial dilutions to minimize carryover. Pure peptides were screened using 3-fold dilution series across 11 concentrations, n=2 times. Fresh pipette tips were used for all serial dilutions.

#### Curve fit, crude peptides

For each peptide sample, the five activity readings were normalized using the corresponding controls. In agonist mode, the positive control was anchored at 0 and the negative control was anchored at 1. For antagonist mode, the normalisation was flipped in each case to always create a downwards sloping curve. Then, the Hill Equation was fitted to the five pairs of concentration and activity using non-linear least squares fitting with constraints. The top parameter and the Hill coefficient were both constrained close to 1. The maximum response parameter was constrained close to 0, corresponding to full efficacy. The potency parameter was never constrained and was always reported as an assay endpoint.

### Biophysics assays for solubility and amyloid fibril formation

Some libraries were assessed for solubility and amyloid fibril formation using turbidity and Thioflavin T (ThT) fluorescence assays. Peptides were dissolved in buffers (50 mM phosphate pH 7.0 and 50 mM acetate pH 4.0) to 267 µM and left 2 hours at room temperature. Two aliquots from each sample (80 µl per well) were transferred to black 384-well microplates with clear bottoms (Greiner Bio-One, Cat. no. 781096) and mixed with ThT (2 µl per well) to a final ThT concentration of 4 µM. The plates were sealed before measurements. Firstly, solubility was evaluated by measuring turbidity as absorbance at 600 nm using a CLARIOstar Plus microplate reader (BMG Labtech). Increased absorbance was interpreted as presence of precipitated material and hence indicative of reduced solubility. For subsequent amyloid fibril formation analysis, the plate was incubated at 40 °C and stressed using alternating cycles of 5 min rest and 5 min linear shaking at 700 rpm over 72 hours within the plate reader. Between cycles fluorescence emission at 480 nm (excitation at 450 nm) was measured. Fibril formation for each sample was determined as the average emission over time.

### Peptide library designs

In vitro maturation of the best de novo hit peptide was done by systematically designing and analysing beneficial substitutions and HLE-conjugations in the DMS, HLE scan, combinatorial scan libraries. Peptides that failed during synthesis were filtered away from the analysis.

#### DMS

The DMS comprised of first, second and triple order peptide variants using all natural amino acids except cysteine and methionine, together with two non-canonical amino acids, norleucine (Nle) and 2-aminoisobutyric acid (Aib). This included all possible single-point substitutions across the sequence (304 peptides) together with a random subset of double and triple substitutions. A total of 835 peptide variants were successfully synthesized and functionally tested for hMC4R agonist and antagonist activity.

#### HLE conjugation scan

In this work, HLE refers to a fatty acid attached to a linker. Octadecanoic acid was denoted C18, and 1,18-octadecanedioic acid denoted C18DA; similarly, eicosanoic acid was denoted C20, and 1,20-eicosanedioic acid, C20DA. For simplicity, C18- and C20-based HLEs were referred to as mono-acids, whereas C18DA- and C20DA-based HLEs referred to as di-acids.

The hit peptide was conjugated with 9 different HLEs at positions 2-16 combined with variations of linker composition, L-γ-glutamyl (γE) and 3,8-dioxa-aminooctanoic acid (OEG) via the sidechain of a lysine substitution (K_ε_): C18-γE, C18-γE-(OEG)_2_, C18-(γE)_2_-(OEG)_2_, C18DA-γE, C18DA-γE-(OEG)_2_, C18DA-(γE)_2_-(OEG)_2_, C20DA-γE, C20DA-γE-(OEG)_2_, C20DA-(γE)_2_-(OEG)_2_. The HLE-conjugated scan comprised of 173 peptides.

#### Combinatorial scan

In the combinatorial library design, we systematically combined the top-performing single-point mutations identified from the DMS with selected amino acids of diverse properties and diacid HLE-conjugation. C18DA-γE-(OEG)2 was used as di-acid because C18 and C20 diacids are among the most commonly used HLEs supporting once-weekly dosing^39^. In total, 181 peptide variants were successfully synthesized and screened in a functional assay.

### Data modelling

Data modelling of the in vitro measurements was performed as described in Nielsen et. al. (2024) with peptide encoding, model prediction, random forest regression and SHAP analysis^31^. Peptide sequences from the DMS were encoded with z-scales^55^ and modelled with a random forest regressor. The predictions from the random forest regression are used in the analysis of positional substitution effects. Peptide sequences from the HLE scan and combinatorial scan are encoded as one-hot-encoded sequences and fitted through a random forest regression, following a SHAP model for investigation of positional impact of substitutions and HLE-conjugations.

Firstly, values outside of dilution window are filtered away and flagged as inactive. To account for systematic batch-to-batch offsets across peptide libraries and experimental runs, potency values in Fig. 2 were batch-corrected by applying an additive shift in log space. Specifically, pEC50 values obtained from crude peptide measurements were corrected using a batch-specific offset derived from the difference between the mean pEC50 of the crude hit peptides measured within each batch and the pEC50 of the corresponding purified hit peptide. Batch correction was performed by:

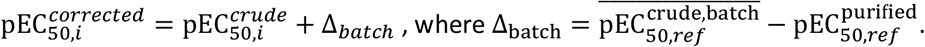

Hit library, DMS and HLE scan libraries are corrected to the best de novo hit peptide, while the combinatorial scan is corrected to the W15KεC18DA variant. We demonstrate that batch correction does not affect rank ordering between crude and purified peptides (Fig. S4). Raw values can be observed in Fig. S5.

### In vivo characterization in mice

#### Animals and housing

Lean male NMRI/RjHan mice were obtained from Janvier (Janvier Laboratories, France), and single-housed in a controlled environment (12/12 h light-dark cycle, lights off at 3 PM, 21 °C ± 2 °C, humidity 50% ± 10%) with ad libitum access to chow (Altromin 1324, Brogaarden, Denmark) and water unless otherwise stated. Mice arrived at 4 weeks of age and were 9 weeks old at study initiation. All procedures were conducted in an AAALAC-accredited facility in accordance with Gubra’s bioethical guidelines and were approved by the Danish Animal Experimentation Council.

#### Intracerebroventricular cannulation

Mice were surgically implanted with a unilateral guide cannula targeting the right lateral ventricle under isoflurane anaesthesia, using stereotaxic coordinates (mouse coordinates (mm) are anterior-posterior: -0.25; medial-lateral: +/-1.0; caudal-ventral: -2.5.). Cannulae were secured with dental cement, and postoperative analgesia was provided for at least 48 hours. Following recovery, compounds were administered via the cannula in awake mice as a single intracerebroventricular bolus injection (5 µL total volume per animal), delivered manually at a maximum flow rate of ∼10 µl min^−1^.

#### Dosing and study design

Mice were fasted for 6 hours prior to dosing, which occurred immediately before the onset of the dark phase (15:00). Animals were randomized by body weight and 24-hour monitoring of food intake one day prior to study start. Animals received a single intracerebroventricular dose of vehicle (50 mM phosphate, 70 mM NaCl, pH 7.4; n = 8) or the E5P peptide variant (10 nmol in a fixed volume of 5μL of a 2000 nmol/mL solution; n = 8).

#### Metabolic phenotyping

Food intake, water intake, and locomotor activity were monitored using an automated metabolic phenotyping system (PhenoMaster, TSE Systems). Locomotor activity was quantified by infrared beam brakes in the X- and Y-axes 64 beams and 32 beams respectively within each cage. Mice were acclimatized to the system prior to dosing. Data were recorded continuously every 10 minutes from 24 hours before dosing and for 24 hours post-dose, with intake and activity quantified automatically without experimenter intervention.

#### Statistical analysis

Single-timepoint continuous endpoints were compared between the vehicle and treatment group using two-tailed unpaired t-test, assuming normally distributed data with homogeneous variance.

## Data availability

Data are available from the corresponding author upon reasonable request.

## Acknowledgements

We thank Mette Elmegaard Andersen, Morten Schlein, Malte Hasle Nielsen, Michele Cavalera, and Henrik Björk Hansen for their assistance with data interpretation and valuable discussions related to this study.

## Author contributions

M.N.Y., J.C.N. and L.S.D. conceptualized the study. M.N.Y. and J.C.N. designed experiments. M.N.Y., V.M., J.M.J. performed de novo design and computational analysis. P.T. carried out peptide synthesis.

R.B.M. and N.B.M. carried out receptor potency assays. A.K. coordinated the in vivo experiments. V.M., J.M.J., M.N.Y., J.C.N., R.B.M., P.T., A.K. and T.P.J. analysed and interpreted data. V.M. and J.M.J. wrote the manuscript. V.M. and J.M.J. contributed equally to this work. All authors reviewed and accepted the paper.

