## Supplementary information for "De novo designed cyclic MC4R peptide agonist reduces food intake in mice"

### Supplementary Figures

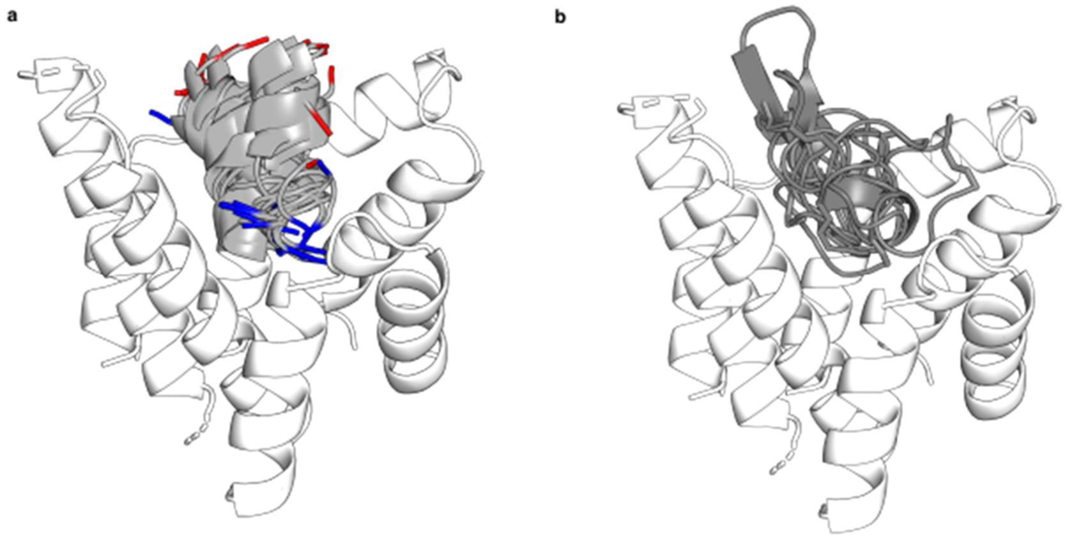


*Fig. S1: Sample of predicted 14 linear and 8 cyclic de novo peptide (dark grey) conformations to the cropped hMC4R structure; PDBID: 6W25 (white). The linear peptides adopt an extended conformation often inserting the N-terminal (blue) in the binding pocket of hMC4R, while C-terminal (red) points away. The cyclic peptides occupy larger volume of the binding pocket.*

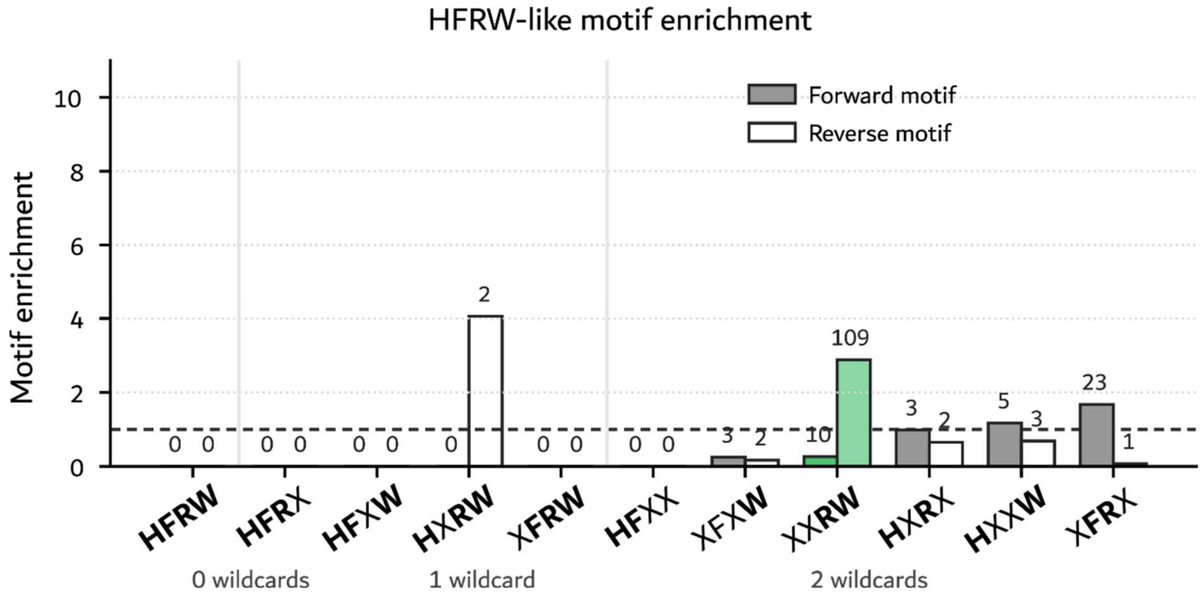


*Fig. S2:* ***Occurrences of known MC4R motifs in cyclic hit library.***
*Forward motifs are shown as* dark *bars, while reversed motifs are shown as* light *bars. Green bars indicate motifs present in the top de novo hit.*

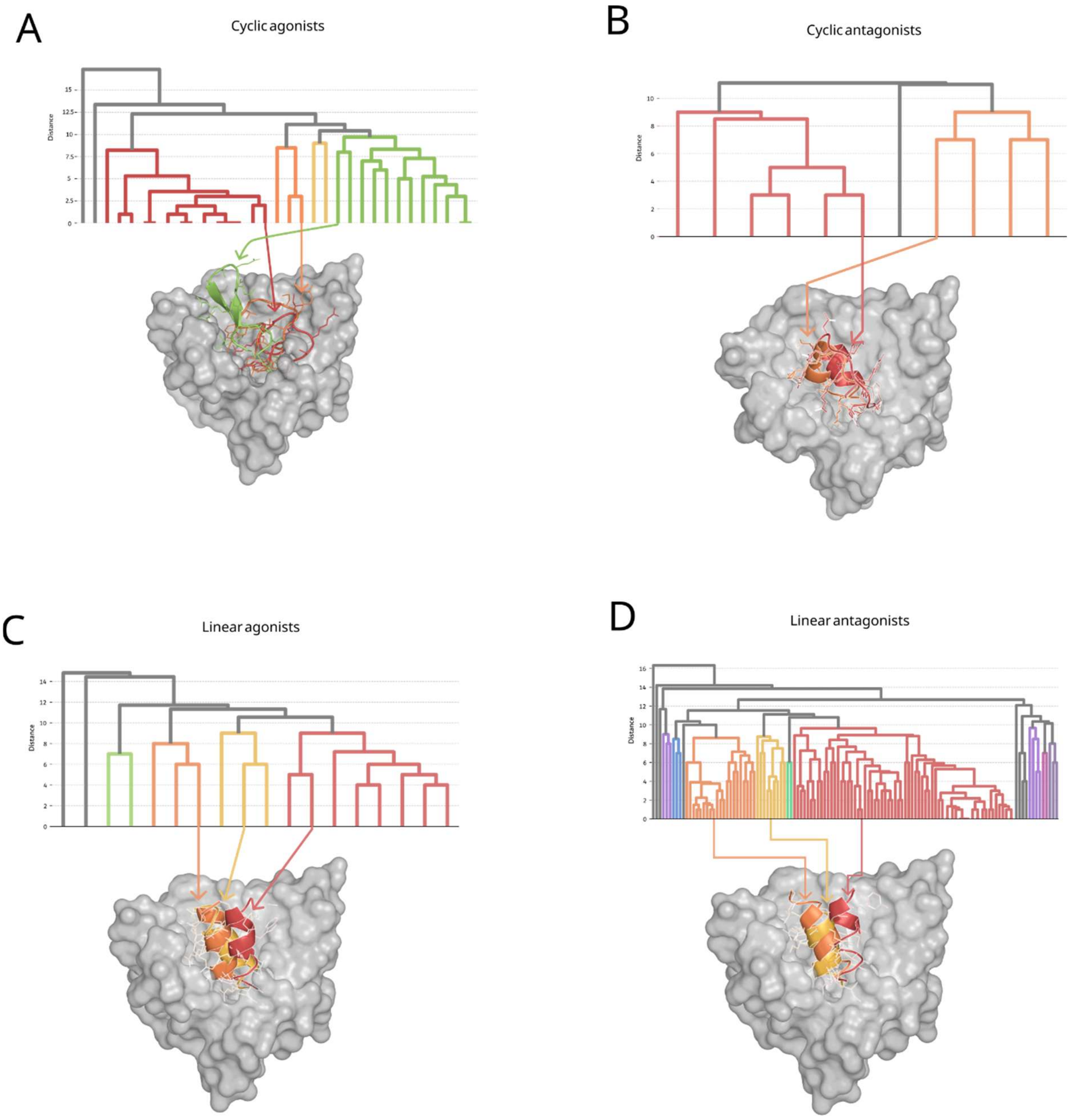


*Fig. S3:* ***Structural diversity of functionally active cyclic and linear de novo designed peptides within the hMC4R binding pocket.***

*Functionally active cyclic and linear peptides were clustered based on Hamming distances between their receptor contact point fingerprints, where each peptide was represented by a binary vector indicating which hMC4R residues were in contact. A contact was defined as any peptide-receptor residue pair with a minimum interatomic distance <4 Å. Clustering was performed using a distance cutoff of 10. All peptides were predicted to bind within the orthosteric binding pocket of hMC4R, but they engage distinct sets of receptor contact points. The dendrograms reveals multiple structural clusters, and representative peptides from three major clusters illustrate the diverse binding conformations adopted within the same pocket.*

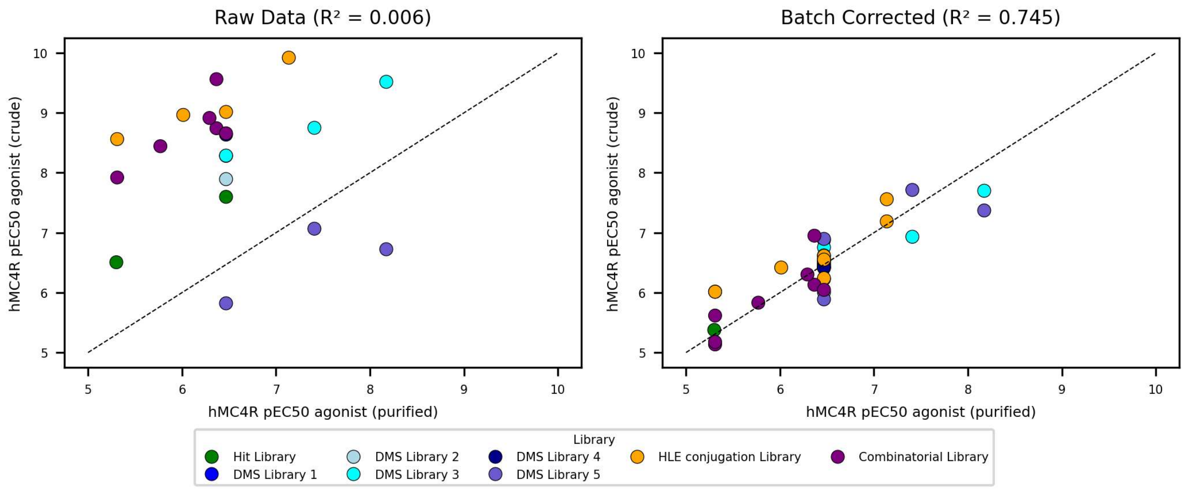


*Fig. S4: Correlation between crude and purified peptides in different libraries on raw crude values before batch correction (left) and after batch correction (right).*

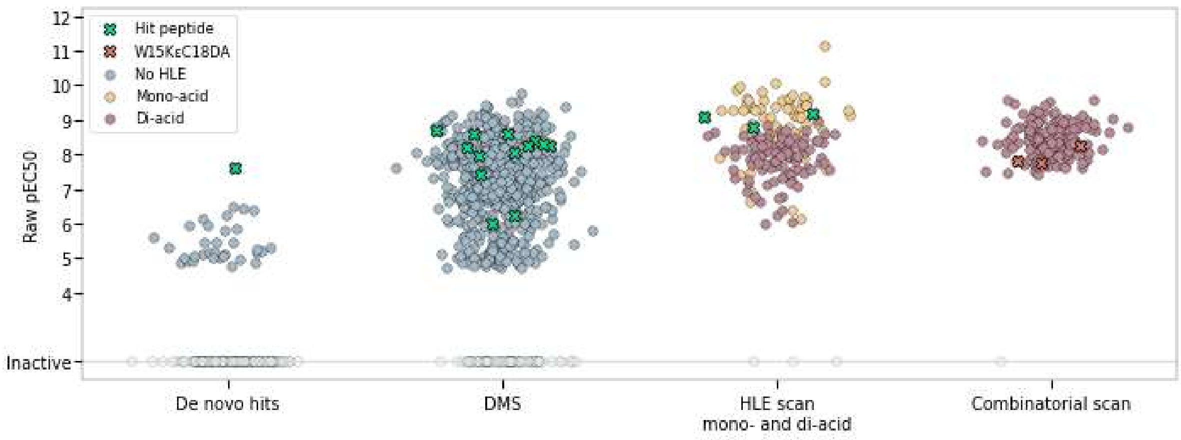


*Fig. S5: Raw pEC50 values of crude peptides in libraries before batch correction.*

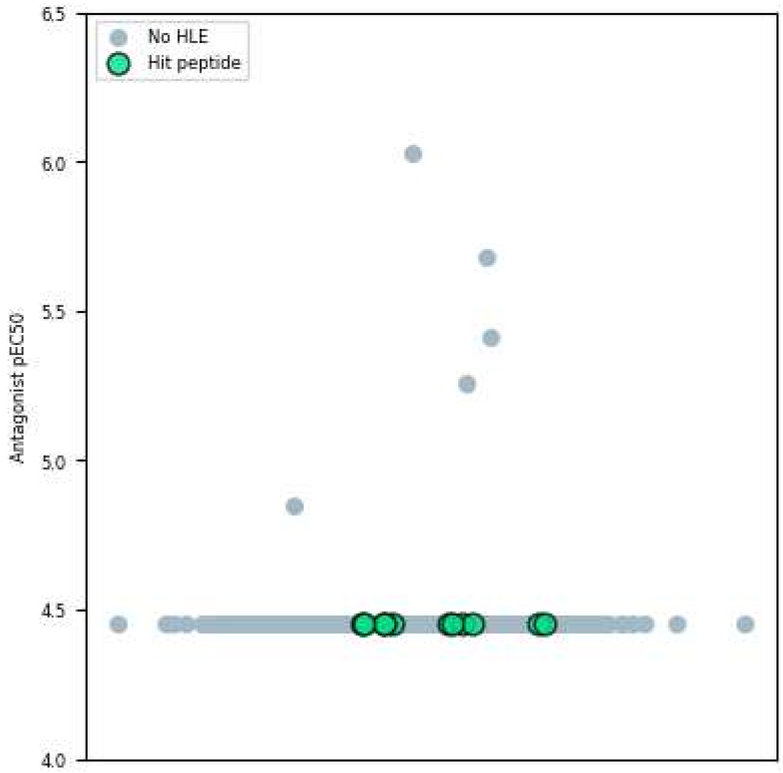


*Fig. S6: hMC4R antagonist pEC50 of DMS compounds*

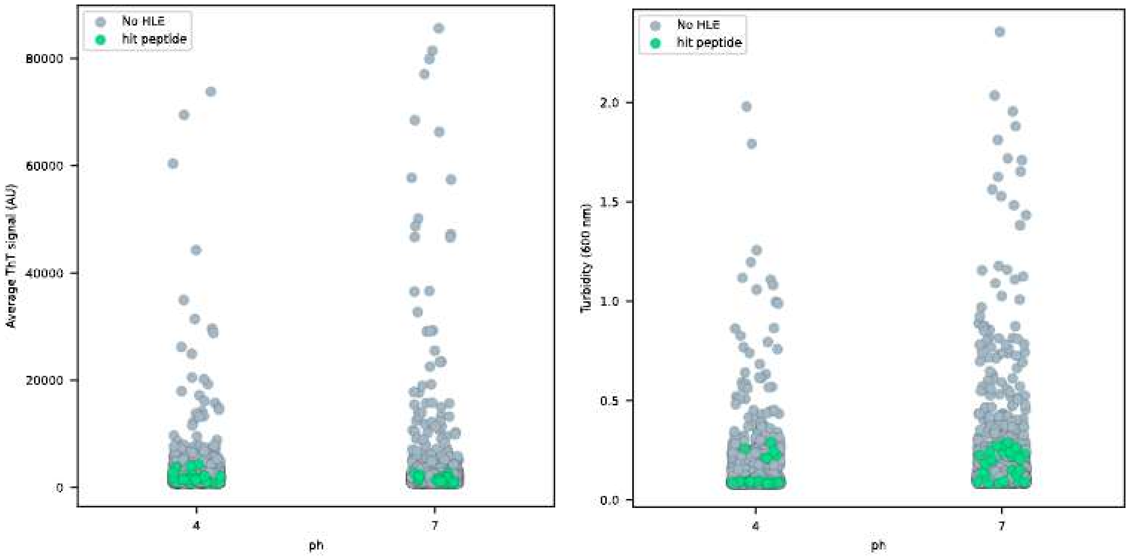


*Fig. S7: Fibril formation (left) and turbidity (right) at pH 4 and 7 of peptides in DMS*

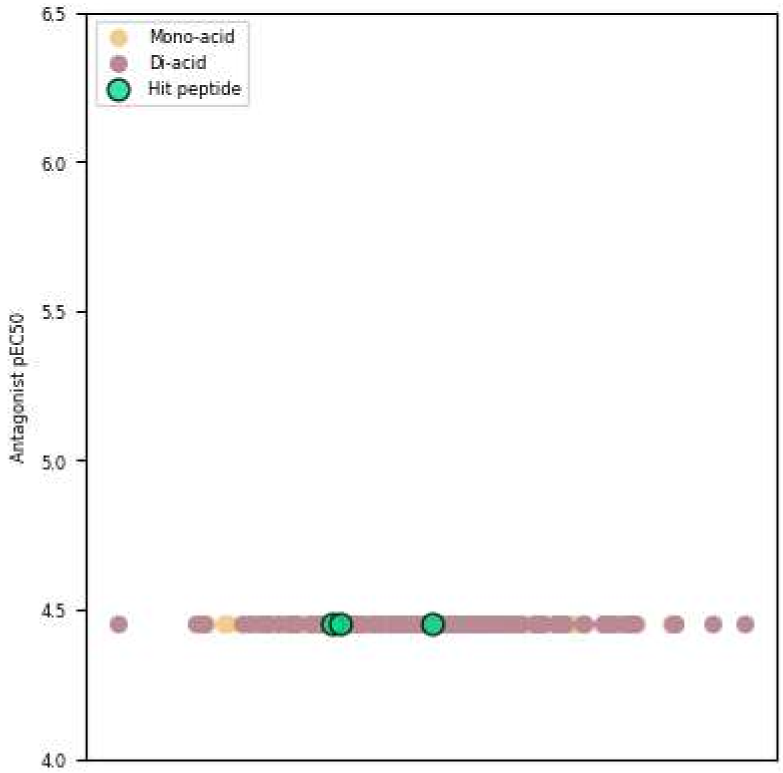


*Fig. S8: hMC4R antagonist pEC50 of peptides in HLE scan*

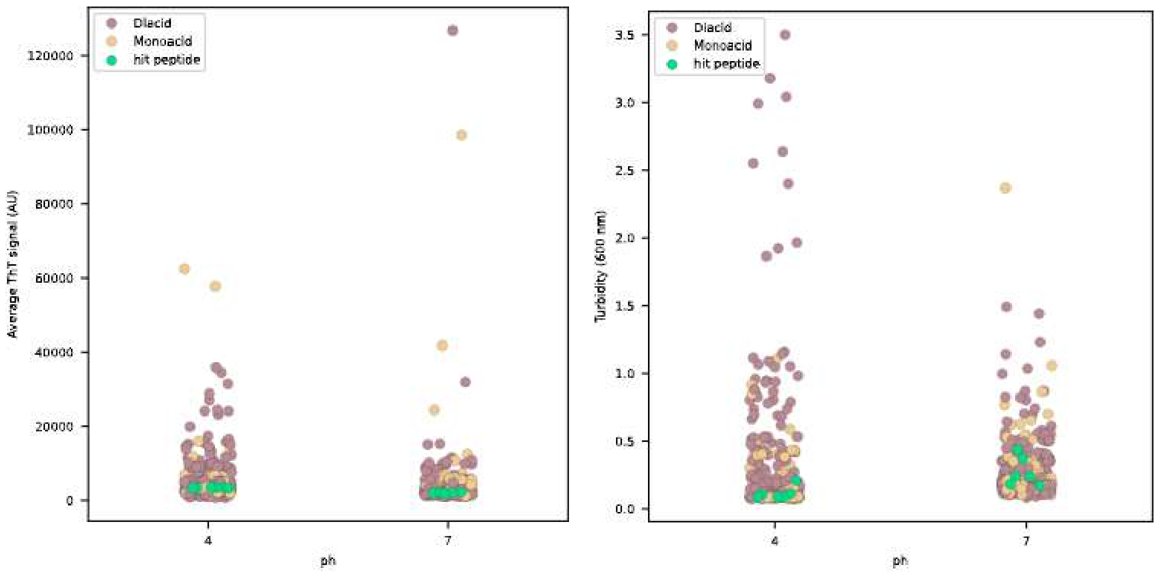


*Fig. S9: Fibril formation (left) and turbidity (right) at pH 4 and 7 of peptides in HLE-conjugation scan*

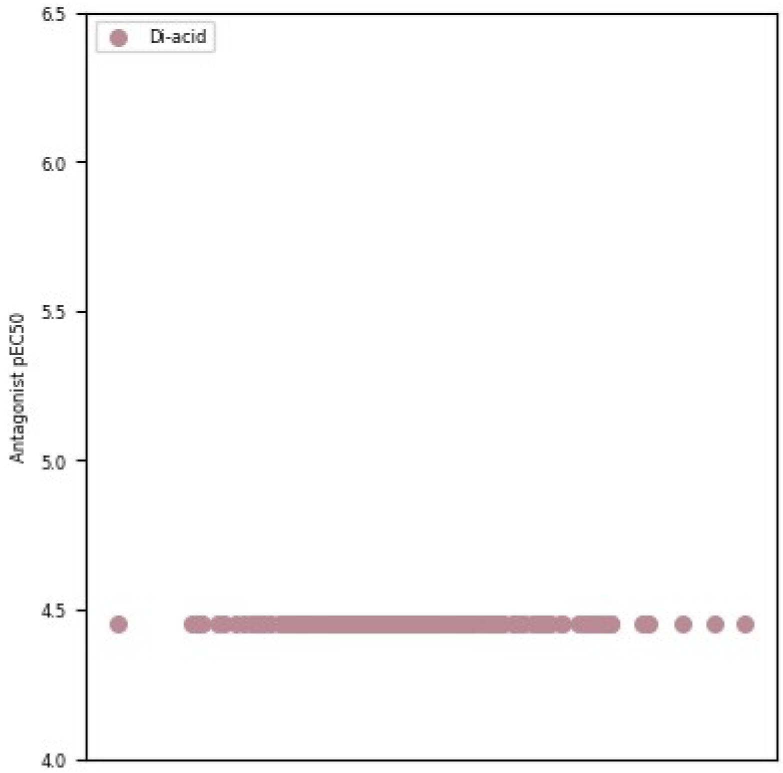


*Fig. S10: hMC4R antagonist pEC50 of peptides in combinatorial scan*

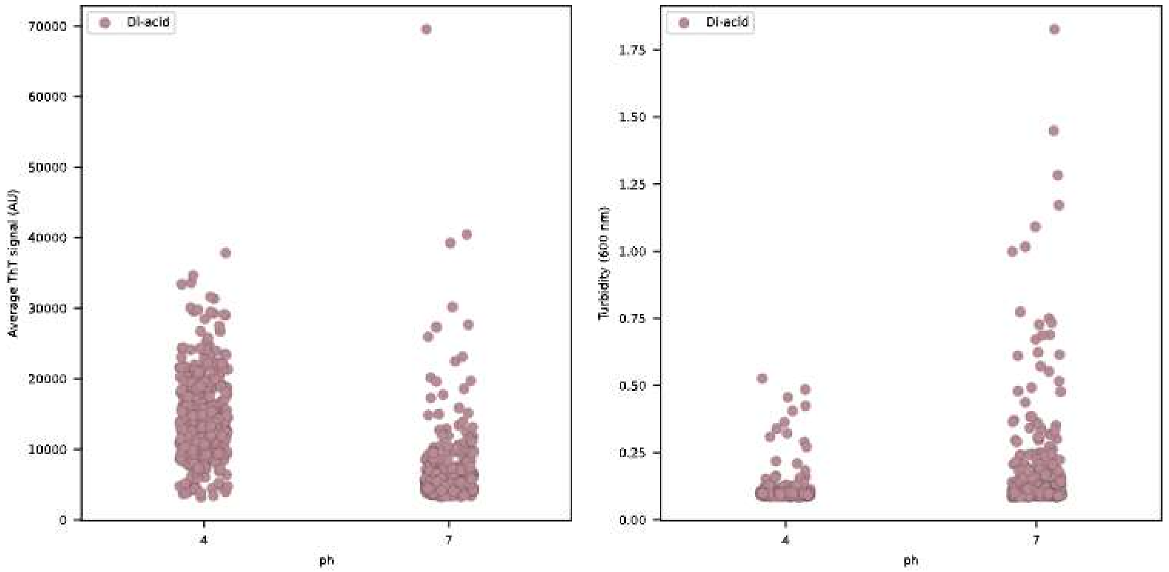


*Fig. S11: Fibril formation (left) and turbidity (right) at pH 4 and 7 of peptides in combinatorial scan*

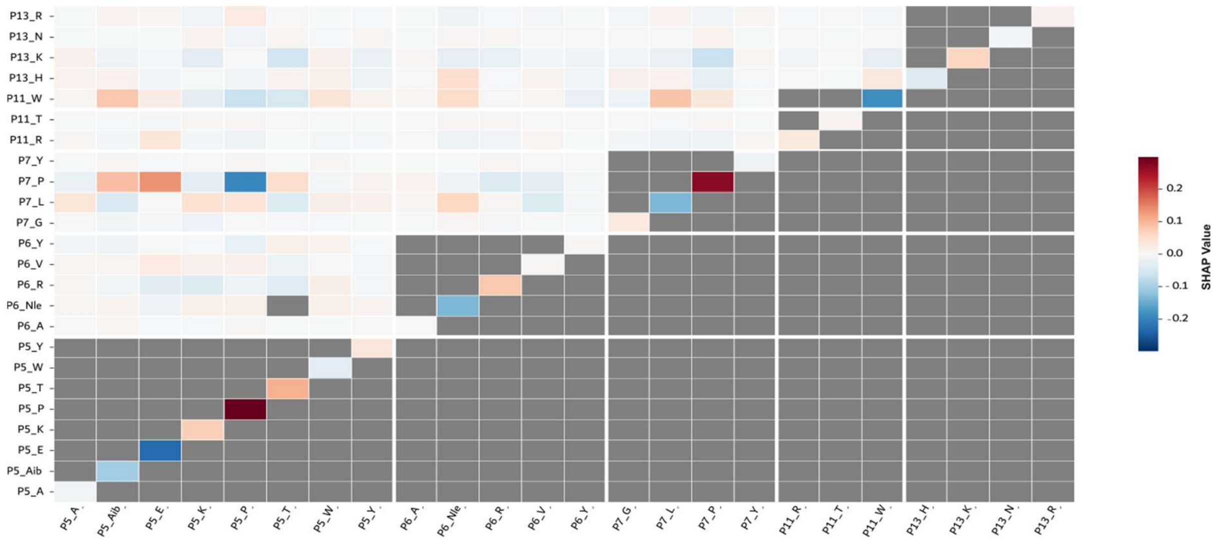


*Fig. S12: SHAP interaction plot of combinatorial design library.*

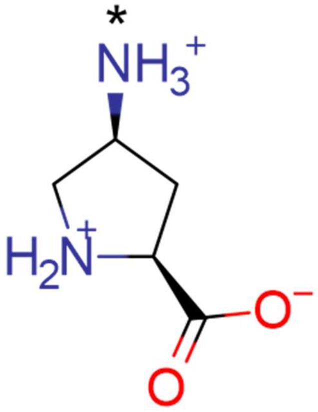

*Fig. S13: Chemical structure of 4-aminoproline ($P_\gamma$). The amine used for HLE conjugation is indicated with an asterisk.*

*Table S1:* ***In vitro cross-reactivity profiling of purified de novo cyclic peptide variants across the melanocortin receptor family.***

*C18 corresponds to the C18-(γE)$_2$-(OEG)$_2$ HLE, and C18DA corresponds to the C18DA-γE-(OEG)$_2$ HLE. HLE conjugations via 4-aminoproline (Pγ). The reported EC50 values are averages of ≥2 independent replicates. Measurements outside the tested concentration range are reported with a “>” in the table (α-MSH and C18DA-based variants: $EC_{50}/IC_{50}$ > 5,000 nM; non-HLE conjugated, C18-based and linear variant: $EC_{50}/IC_{50}$ > 50,000 nM). Non-measured potencies are annotated with N/D.*

| Variant | Category | hMC4R | | hMC1R | | hMC2R | | hMC3R | | hMC5R | |
|---|---|---|---|---|---|---|---|---|---|---|---|
| | | *Agonism* | *Antagonism* | *Agonism* | *Antagonism* | *Agonism* | *Antagonism* | *Agonism* | *Antagonism* | *Agonism* | *Antagonism* |
| | | *EC50 [nM]* | *IC50 [nM]* | *EC50 [nM]* | *IC50 [nM]* | *EC50 [nM]* | *IC50 [nM]* | *EC50 [nM]* | *IC50 [nM]* | *EC50 [nM]* | *IC50 [nM]* |
| α-MSH | No HLE | 0.39 | >5000 | 0.14 | >5000 | >5000 | >5000 | 0.47 | >5000 | 5.0 | >5000 |
| Hit peptide | | 340 | >50,000 | 450 (61%) | >50,000 | >50,000 | >50,000 | >50,000 | 500 | >50,000 | 760 |
| E5P | | 6.7 | >50,000 | 73 | >50,000 | >50,000 | >10,000 | 210 (38%) | N/D | 29 | >50,000 |
| Y7P | | 39 | >50,000 | 580 (74%) | >50,000 | >50,000 | >50,000 | >50,000 | 1600 | 690 | >50,000 |

| | | | | | | | | | | | |
|---|---|---|---|---|---|---|---|---|---|---|---|
| W15KεC18 | C18 | 73 | >50,000 | 31 (59%) | >10,000 | >50,000 | 210 | >50,000 | 61 | >50,000 | 250 |
| Y7P:W15KεC18 | | 55 | >50,000 | 43 (59%) | >50,000 | >50,000 | 210 | >50,000 | 41 | >50,000 | 210 |
| E5KεC18:Y7P | | 27 | >50,000 | 8.6 | >50,000 | >50,000 | 90 | >50,000 | 1.9 | >50,000 | 24 |
| W15KεC18DA | C18DA | 4900 | >5000 | >5000 | >5000 | >5000 | >5000 | >5000 | >5000 | >5000 | >5000 |
| E5P:W15KεC18DA | | 430 | >5000 | 1000 | >5000 | >5000 | >5000 | >5000 | >5000 | >5000 | >5000 |
| Y7P:W15KεC18DA | | 340 | >5000 | 960 | >5000 | >5000 | >5000 | >5000 | >5000 | >5000 | >5000 |
| E5T:W15KεC18DA | | 1700 | >5000 | 1000 | >5000 | >5000 | >5000 | >5000 | >5000 | >5000 | >5000 |
| E5T:Y7P:W15KεC18DA | | 430 | >5000 | 340 | >5000 | >5000 | >5000 | >5000 | >5000 | >5000 | >5000 |
| E5P:Y7P:W15KεC18DA | | 510 | >5000 | 990 | >5000 | >5000 | >5000 | >5000 | >5000 | >5000 | >5000 |
| E5KεC18DA | | 970 | N/D | N/D | N/D | N/D | N/D | N/D | N/D | N/D | N/D |
| E5KεC18DA:Y7P | | 480 | N/D | N/D | N/D | N/D | N/D | N/D | N/D | N/D | N/D |
| Y7PγC18DA | | 300 | N/D | N/D | N/D | N/D | N/D | N/D | N/D | N/D | N/D |
| E5PγC18DA | | 360 | N/D | N/D | N/D | N/D | N/D | N/D | N/D | N/D | N/D |
| A1PγC18DA | | >5000 | N/D | N/D | N/D | N/D | N/D | N/D | N/D | N/D | N/D |
| A1KεC18DA | | >5000 | N/D | N/D | N/D | N/D | N/D | N/D | N/D | N/D | N/D |
| A1KεC18DA:E5P | | >5000 | N/D | N/D | N/D | N/D | N/D | N/D | N/D | N/D | N/D |
| F16FNH2 | Linear | >50,000 | N/D | N/D | N/D | N/D | N/D | N/D | N/D | N/D | N/D |
| Y7P:W15KεC18DA:F16FNH2 | | >50,000 | N/D | N/D | N/D | N/D | N/D | N/D | N/D | N/D | N/D |
| E5T:Y7P:W15KεC18DA:F16FNH2 | | >50,000 | N/D | N/D | N/D | N/D | N/D | N/D | N/D | N/D | N/D |

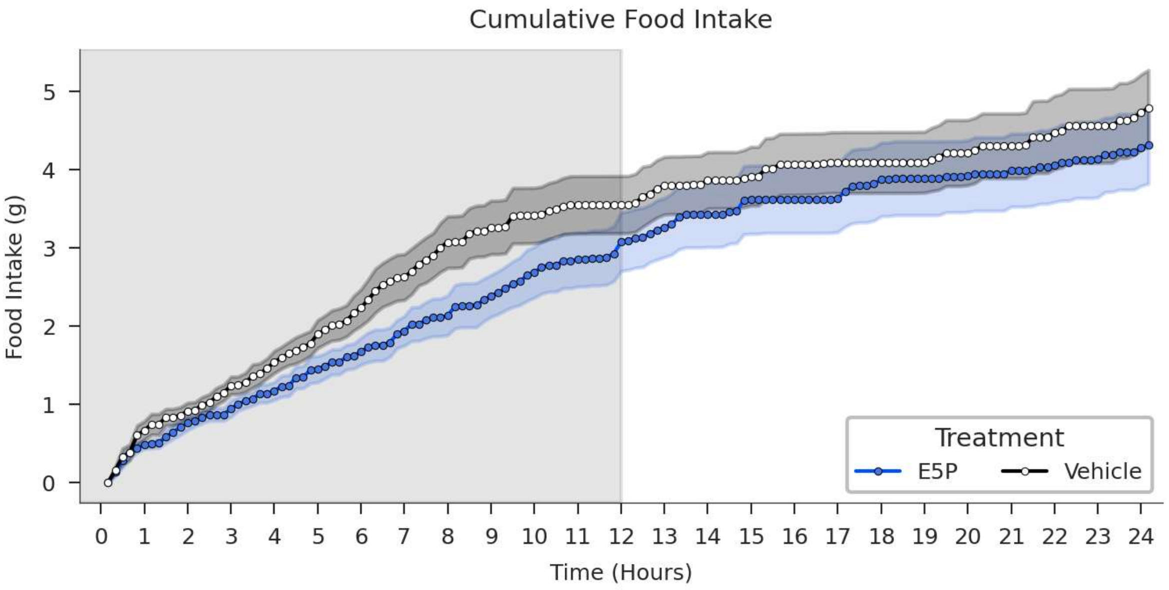


*Fig. S14: Cumulative food (upper) and water (lower) intake over 24 hours of ICV-dosed animals between vehicle-treated (white) and E5P-treated (blue) mice. Shaded area around curves corresponds to average ± SEM. The left grey boxes indicate dark phase.*

*Table S2: **Weight and food intake of treatment groups prior to dosing**.*
*Body weight and 24-hour food intake of animals 1 day prior to dosing between treatment groups. The average ± standard deviation of the weight prior to dosing is 38.11±1.94 and 38.13±2.04 for the vehicle and E5P-treated group, respectively. The average ± standard deviation of the 24-hour food intake prior to dosing is 5.05±0.42 and 5.06±0.47 for the vehicle and E5P-treated group, respectively*

| Animal | Treatment | Weight prior to dosing (g) | Food intake (g) |
|---|---|---|---|
| 1 | Vehicle | 37.80 | 4.90 |
| 2 | Vehicle | 38.00 | 4.51 |
| 3 | Vehicle | 38.40 | 5.32 |
| 4 | Vehicle | 37.70 | 4.65 |
| 5 | Vehicle | 34.20 | 4.82 |

| | | | |
|---|---|---|---|
| 6 | Vehicle | 38.00 | 4.87 |
| 7 | Vehicle | 39.00 | 5.57 |
| 8 | Vehicle | 41.80 | 5.78 |
| 9 | E5P | 37.90 | 5.31 |
| 10 | E5P | 35.80 | 4.71 |
| 11 | E5P | 37.10 | 4.35 |
| 12 | E5P | 37.80 | 5.59 |
| 13 | E5P | 37.40 | 4.83 |
| 14 | E5P | 43.20 | 4.61 |
| 15 | E5P | 38.20 | 5.30 |
| 16 | E5P | 37.60 | 5.75 |

*Table S3:* ***Statistical comparison of cumulative intake between treatment groups***.
*Cumulative food and water intake in vehicle- and E5P-treated mice at 1, 4, 8, and 24 hours post-dosing. Statistical analysis was performed using two-tailed unpaired t-test against vehicle group. Reported p-values reflect comparisons between treatment groups at each time point.*

| Treatment | Hour | Number | Average | SD | SEM | p-value |
|---|---|---|---|---|---|---|
| **Food** | | | | | | |
| Vehicle | 1 | 8 | 0.664 | 0.329 | 0.116 | |
| E5P | 1 | 8 | 0.479 | 0.147 | 0.052 | 0.169 |
| Vehicle | 4 | 8 | 1.54 | 0.404 | 0.143 | |
| E5P | 4 | 8 | 1.17 | 0.322 | 0.114 | 0.0617 |
| Vehicle | 8 | 8 | 3.07 | 0.933 | 0.33 | |
| E5P | 8 | 8 | 2.14 | 0.715 | 0.253 | 0.0408 |
| Vehicle | 24 | 8 | 4.74 | 1.37 | 0.484 | |
| E5P | 24 | 8 | 4.28 | 1.38 | 0.488 | 0.52 |
| **Drink** | | | | | | |
| Vehicle | 1 | 8 | 0.642 | 0.57 | 0.202 | |
| E5P | 1 | 8 | 0.151 | 0.182 | 0.0645 | 0.0359 |
| Vehicle | 4 | 8 | 1.69 | 0.883 | 0.312 | |
| E5P | 4 | 8 | 0.599 | 0.592 | 0.209 | 0.0119 |
| Vehicle | 8 | 8 | 3.13 | 1.61 | 0.569 | |
| E5P | 8 | 8 | 1.6 | 0.986 | 0.349 | 0.038 |
| Vehicle | 24 | 8 | 4.63 | 1.8 | 0.636 | |
| E5P | 24 | 8 | 4.07 | 1.43 | 0.506 | 0.504 |
